# Alteration of Water Exchange Rates Following Focused Ultrasound-Mediated BBB Opening in the Dorsal Striatum of Non-Human Primates: A Diffusion-Prepared pCASL Study

**DOI:** 10.1101/2025.07.29.667530

**Authors:** Dong Liu, Xingfeng Shao, Fabian Munoz Silva, Soroosh Sanatkhani, Ray Lee, Elisa E Konofagou, Danny JJ Wang, Vincent P Ferrera

## Abstract

This study applied diffusion-prepared pseudo-continuous arterial spin labeling (DP-pCASL) to quantify cerebral blood flow (CBF), arterial transit time (ATT), and blood-brain barrier (BBB) water exchange rate (*K*_*w*_) before and after focused ultrasound (FUS)-mediated blood-brain barrier opening (BBBO) in the dorsal striatum of four non-human primates. Six baseline and seven BBBO sessions were performed. DP-pCASL was acquired approximately 45 minutes after FUS sonication combined with intravenous microbubbles, and contrast-enhanced T1-weighted imaging was subsequently used to confirm the BBBO region. Whole-brain analyses revealed no significant changes in CBF or ATT following BBBO (permutation p > 0.05). Region-of-interest analysis within the sonicated caudate demonstrated a significant localized decrease in *K*_*w*_, with median (IQR) values of 45.0 (40.6 - 55.6) min^−1^ at the BBBO site versus 61.6 (58.3 - 70.4) min^−1^ in the contralateral control region (p < 0.05), confirming spatially specific suppression of transendothelial water flux. In contrast, whole-brain *K*_*w*_ increased significantly following BBBO, with median (IQR) values of 49.8 (46.3 - 55.9) min^−1^ in non-BBBO sessions versus 59.4 (56.6 - 66.3) min^−1^ in BBBO sessions (p < 0.01), indicating a diffuse enhancement of water exchange across the brain. These findings establish DP-pCASL-derived *K*_*w*_ as a sensitive, non-contrast biomarker for both local and global BBB permeability changes induced by focused ultrasound, supporting its potential for longitudinal monitoring in preclinical and clinical neurotherapeutic applications.

## Introduction

Focused ultrasound (FUS) combined with intravenously administered microbubbles enables a safe, transient, and localized opening of the blood-brain barrier (BBB) (Aryal et al., 2014; Hynynen et al., 2001; McDannold et al., 2007). This effect is achieved by applying focused acoustic energy that causes circulating microbubbles within targeted cerebral vessels to oscillate (Marquet et al., 2014; Tung et al., 2011), generating mechanical forces that temporarily disrupt endothelial tight junctions (Konofagou, 2012; Sheikov et al., 2004). By temporarily disrupting the BBB, FUS facilitates the delivery of therapeutic agents such as chemotherapeutic drugs, antibodies, and viral vectors into selected brain regions, offering promising treatments for neurological and psychiatric disorders including Alzheimer’s disease, Parkinson’s disease, and brain cancers (Leinenga et al., 2016; Lipsman et al., 2018; Mainprize et al., 2019; Rezai et al., 2020). While these agents primarily penetrate the brain via paracellular diffusion through the disrupted tight junctions, additional pathways such as receptor-mediated transcytosis and adsorptive endocytosis may also facilitate their transport (Burgess et al., 2015; Hawkins and Davis, 2005). Similarly, gadolinium-based contrast agents, which normally cannot cross an intact BBB, gain access through these temporary junctional openings and accumulate in brain tissue. This characteristic underpins the use of gadolinium-enhanced magnetic resonance imaging (MRI) as the standard method for detecting and confirming BBB opening in both research and clinical settings (McDannold et al., 2006). However, concerns regarding the safety of repeated gadolinium administration, including risks of nephrogenic systemic fibrosis and gadolinium retention in brain tissue, have prompted the development of contrast-free imaging methods to safely monitor BBB integrity over repeated treatments (Gulani et al., 2017; Kanda et al., 2014).

In addition to facilitating molecular delivery, focused ultrasound-induced BBB opening (FUS-BBBO) can modulate neural activity, triggering significant changes in brain functional connectivity and neuroinflammation, as well as influencing behavior outcomes. For instance, studies in non-human primates have reported enhanced cognitive performance, including improved response speed and accuracy, following targeted striatal FUS-BBBO, indicating neuromodulatory effects beyond simple molecular delivery (Chu et al., 2015; Pouliopoulos et al., 2021; Todd et al., 2019). Functional MRI studies have demonstrated that FUS-BBBO can alter resting-state functional connectivity in both animals and humans, suggesting transient disruption of network dynamics (Meng et al., 2019; Todd et al., 2018). Notably, a recent non-human primate study compared the effects of FUS-neuromodulation and FUS-BBBO, showing that BBB opening alone can induce robust, region-specific changes in connectivity, further supporting a neuromodulatory role that does not depend on drug delivery (Liu et al., 2023). In parallel, FUS-BBBO has been shown to elicit transient neuroinflammatory responses, including glial activation, which may further contribute to downstream plasticity and behavioral effects (Kovacs et al., 2017; McMahon and Hynynen, 2017). Despite accumulating evidence of these neuromodulatory phenomena, the precise physiological mechanisms underlying these broader effects remain incompletely understood, necessitating deeper exploration into the cellular, molecular, and functional changes triggered by transient FUS-BBBO.

A key factor in maintaining neurovascular homeostasis, which may potentially be involved in the neuromodulatory effects of FUS-BBBO, is the regulation of water transport across the BBB. Aquaporin-4 (AQP4), a water channel protein abundantly expressed on astrocytic endfeet at the gliovascular interface, facilitates water exchange in response to osmotic and hydrostatic gradients, modulating cerebral edema, neuroinflammation, and glymphatic clearance (Papadopoulos and Verkman, 2013). Given its small molecular size (∼18 Da), water acts as a highly sensitive endogenous tracer, capable of detecting subtle permeability alterations that larger molecules such as gadolinium (∼550 Da) might fail to reveal (Dickie et al., 2020; Shao et al., 2020; Uchida et al., 2023). Thus, studying water transport dynamics offers an exceptionally sensitive and physiologically relevant avenue to detect and better understand subtle BBB permeability and functional changes induced by FUS-BBBO.

Diffusion-prepared pseudo-continuous arterial spin labeling (DP-pCASL) is a recently developed, non-invasive MRI technique to quantify the water exchange rate (*K*_*w*_) across the BBB. By combining diffusion weighting with ASL and employing multi-post-labeling delay acquisitions, DP-pCASL improves *K*_*w*_ accuracy by accounting for arterial transit time (ATT) variability (Shao et al., 2019; St. Lawrence et al., 2012; Wang et al., 2007). Clinical studies have reported changes of BBB *K*_*w*_ in association with aging (Shao et al., 2024) and various neurological conditions, including multiple sclerosis (Wengler et al., 2020), obstructive sleep apnea (Palomares et al., 2015) and cerebral small vessel disease (Li et al., 2023). In Alzheimer’s disease, Chen et al. reported significantly reduced *K*_*w*_ in the hippocampus, cingulate cortex, and other brain regions, with associations to plasma biomarkers and cognitive performance, suggesting *K*_*w*_ as a promising biomarker for early BBB dysfunction (Chen et al., 2025). Additionally, Tiwari et al. demonstrated the sensitivity of diffusion-weighted ASL MRI to mannitol-induced BBB disruption in rodents, validated by contrast-enhanced MRI and histology (Tiwari et al., 2025).

Despite these methodological advancements, DP-pCASL has not yet been applied to characterize the changes in water exchange dynamics associated with FUS-induced BBB opening. The aim of the present study, therefore, is to quantify alterations in cerebral blood flow (CBF) and BBB water exchange rate (*K*_*w*_) following FUS-BBBO targeting the dorsal striatum of non-human primates (NHPs) (**Figure 1**). By directly comparing these gadolinium-free DP-pCASL measures with conventional gadolinium-enhanced MRI, we seek to evaluate the potential of DP-pCASL as a non-contrast imaging approach for assessing BBB permeability and to gain insight into the mechanisms by which focused ultrasound transiently alters BBB integrity.

**Figure 1.**
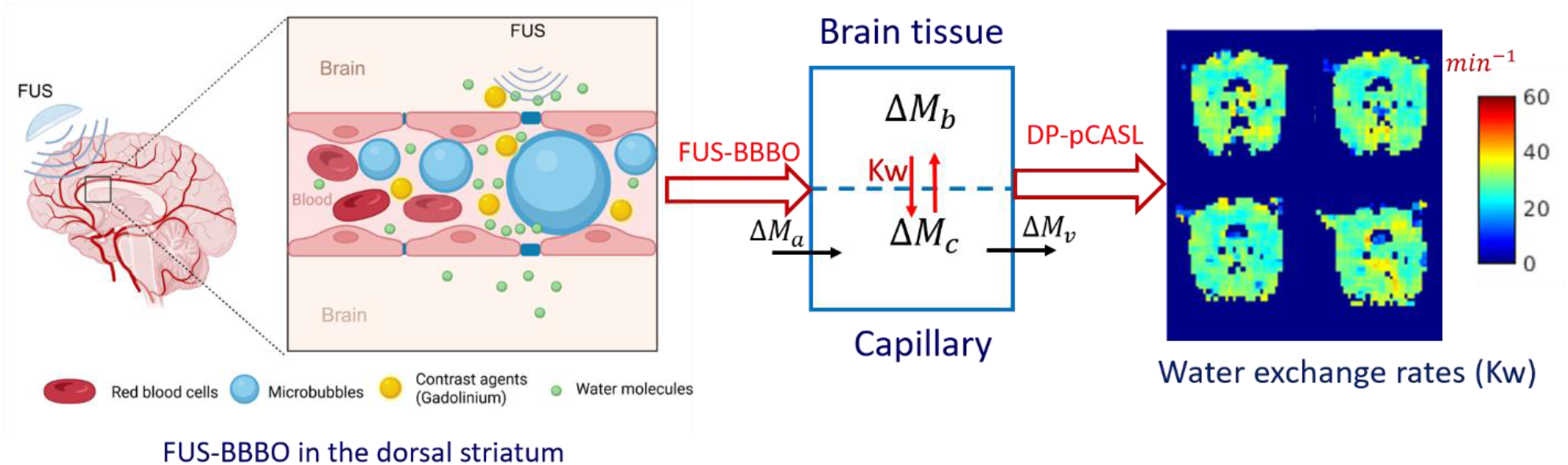
Schematic pipeline for measuring blood–brain barrier (BBB) water exchange between capillary and brain tissue using diffusion-prepared pseudo-continuous arterial spin labeling (DP-pCASL) after focused ultrasound (FUS)–mediated BBB opening in non-human primates (NHPs). Part of the figure was created using BioRender.com. Δ*M*_*a*_, Δ*M*_*b*_, Δ*M*_*c*_ and Δ*M*_*v*_ represent the magnetization of labeled water in the arterial, brain tissue, capillary, and venous compartments, respectively. *K*_*w*_ denotes the water exchange rate across BBB.

## Materials and Methods

### Animal Preparation

Four adult male NHPs, 2 long-tailed macaques (*Macaca fascicularis*) and 2 rhesus macaques (*Macaca mulatta*), weighing between 7.1 and 14.1 kg, were included in this study. Prior to the ultrasound and MRI procedures, animals were initially sedated with ketamine (10 mg/kg) and dexmedetomidine (0.02 mg/kg). Anesthesia was then maintained with inhaled isoflurane at concentrations ranging from 0.8% to 1.1% throughout the experiments. The scalp hair was shaved, and a conductive gel was applied to optimize acoustic coupling for ultrasound delivery. During MRI scanning, physiological parameters—including body temperature, electrocardiogram, oxygen saturation, and end-tidal CO_2_—were continuously monitored using a wireless MRI-compatible system (Iradimed 3880). To minimize potential confounding effects from repeated anesthesia and to ensure complete restoration of BBB integrity between experiments, a strict wash-out period was enforced. Specifically, the interval between consecutive MRI sessions for each subject was maintained at a minimum of 10 days. This duration significantly exceeds the typical 48–72 hour window required for BBB closure and functional recovery in non-human primates (Marquet et al., 2014; Pouliopoulos et al., 2021). Consequently, the frequency of scanning was limited to no more than two sessions per month per animal. All experimental protocols were conducted in accordance with guidelines approved by the Institutional Animal Care and Use Committee (IACUC) of Columbia University.

### FUS with microbubbles application

A single-element FUS transducer (model H-107; 500 kHz frequency; radius of curvature 63.2 mm; outer diameter 64 mm; Sonic Concepts, Bothell, WA) was used for all sonication. The transducer was driven by a function generator (Agilent 33220A, Agilent Technologies, Santa Clara, CA) connected to a 57 dB radiofrequency power amplifier (500S06, E&I, Rochester, NY). The transducer’s acoustic output was calibrated in a degassed water tank using a capsule hydrophone (HGL-0200, Onda Corp., Sunnyvale, CA), which measured free-field pressure, focal location, and acoustic intensities including spatial-peak pulse average (I_sppa_) and spatial-peak temporal average (I_spta_). To estimate the actual acoustic pressure reaching the brain, attenuation through *ex vivo* NHP skulls was measured. To estimate the in situ acoustic pressure, we considered a transcranial attenuation range of 60–65% due to the skull, a value consistent with previous numerical and experimental characterizations (T. Deffieux and E. E. Konofagou, 2010; Wu et al., 2018). In this study, we adopted a conservative 65% reduction to report the derated peak negative pressure (PNP), thereby accounting for potential losses due to skull heterogeneity and incidence angles.

On the day of MRI scanning, the transducer cone was filled with degassed water using a WDS105+ water degassing system (Sonic Concepts, Bothell, WA) and attached securely to the NHP’s scalp to optimize ultrasound transmission. Sonication parameters were based on established protocols to safely and effectively induce BBB opening in NHPs. Animals received sonication for 2 minutes at a derated PNP of 400 kPa, with a pulse repetition frequency (PRF) of 2 Hz, pulse duration of 10 ms, and a 2% duty cycle, as illustrated in **Figure 2**. In-house manufactured microbubbles (lipid-shelled with a perfluorobutane core; diameter 4–5 μm; concentration 2.5 × 10^8^ bubbles/kg) were prepared and stored at 4°C following established protocols (Feshitan et al., 2009; Tung et al., 2011). On the day of the experiment, microbubbles were activated via mechanical agitation immediately prior to use. They were administered as a bolus injection intravenously via the saphenous vein approximately 10 seconds after the start of sonication (injection duration < 10 s). The spatial peak temporal average intensity (Ispta) for BBB opening was 39.2 mW/cm^2^. This ultrasound intensity was selected to reliably open the BBB while maintaining safety, as supported by prior studies(Liu et al., 2023; Pouliopoulos et al., 2021).

**Figure 2.**
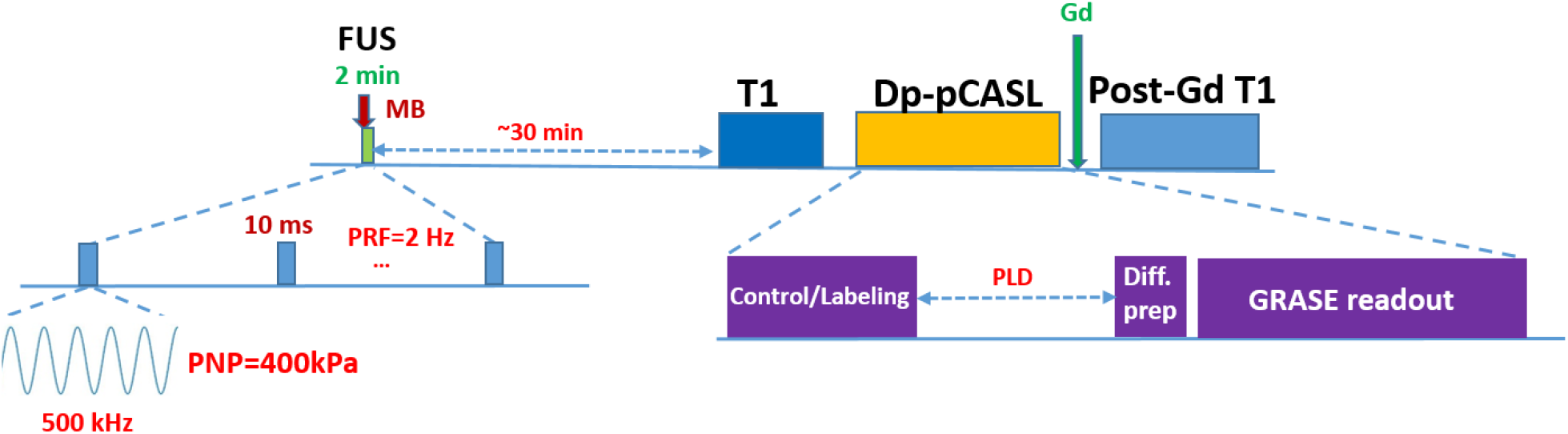
FUS procedures and MRI acquisition: A 2-min focused ultrasound (FUS) exposure combined with microbubbles (MB) was followed by MRI scans, including T1, diffusion-prepared pCASL (2.2 × 2.2 × 5 mm^3^ resolution, 600-1700 ms post-labeling delays (PLDs), with diffusion weighted), and a post-Gadolinium T1 scan to confirm blood-brain barrier (BBB) opening.

### MRI Acquisition

MRI data were acquired on a 3T Siemens Prisma scanner using a 15-channel knee coil optimized for imaging the NHP brain, and the detailed MRI timeline is shown in **Figure 2**. Structural T1-weighted images were obtained using a T1-MPRAGE sequence with the following parameters: repetition time (TR) = 2580 ms, echo time (TE) = 2.81 ms, inversion time (TI) = 1160 ms, flip angle (FA) = 9°, isotropic resolution of 0.5 mm^3^, and field of view (FOV) = 128 × 128 × 120 mm^3^.

DP-pCASL with background-suppressed 3D gradient-and-spin-echo (GRASE) readout was performed within 45 minutes after FUS sonication to measure ATT, CBF, and the BBB *K*_*w*_, following a protocol similar to that described by Shao et al. (Shao et al., 2019). The detailed parameters included: field of view (FOV) = 138 mm, matrix size = 64 × 64, 14 axial slices, voxel size = 2.2 × 2.2 × 5 mm^3^, turbo factor = 14, EPI factor = 64, bandwidth = 2232 Hz/pixel, TE = 31.84 ms, TR = 3200 ms, labeling/control duration = 1000 ms, labeling offset = 55 mm, and centric k-space ordering. Multiple post-labeling delays (PLDs) (600, 1400 and 1700 ms) were acquired with diffusion weightings (b-values) applied to capture water exchange dynamics.

To confirm FUS-mediated BBBO, T1-weighted images were acquired using a T1-SPACE protocol, selected for its sensitivity to low gadolinium concentrations (Danieli et al., 2019), within 20 minutes following intravenous administration of gadolinium-based contrast agent at a dose of 0.2 mmol/kg. The T1-SPACE images matched the pre-contrast T1 and resolution to enable direct comparison.

### MR Image Analysis

A two-stage estimation approach was used to quantify CBF, ATT and BBB *K*_*w*_ from perfusion images acquired at specific post-labeling delays (PLD) and diffusion weightings (b-values) (St. Lawrence et al., 2012), as described in **Figure 3**. In the first stage, ATT was estimated using the flow-encoding arterial spin tagging (FEAST) method, calculated from the ratio of vascular-suppressed signals (with diffusion weighting = 10 s/mm^2^ and velocity encoding [VENC] = 7.5 mm/s) to the total perfusion signals acquired at a short PLD of 600 ms (Wang et al., 2003). In the second stage, *K*_*w*_ was calculated by fitting data acquired at two longer PLDs of 1,400 ms and 1,700 ms with diffusion weightings of b = 0 and 25 s/mm^2^, which allows separation of vascular and extravascular compartments using a two-compartment single-pass approximation (SPA) model (Shao et al., 2019; St. Lawrence et al., 2012). CBF was computed from perfusion signals measured at PLD = 1,700 ms without diffusion weighting (Alsop et al., 2015).

**Figure 3.**
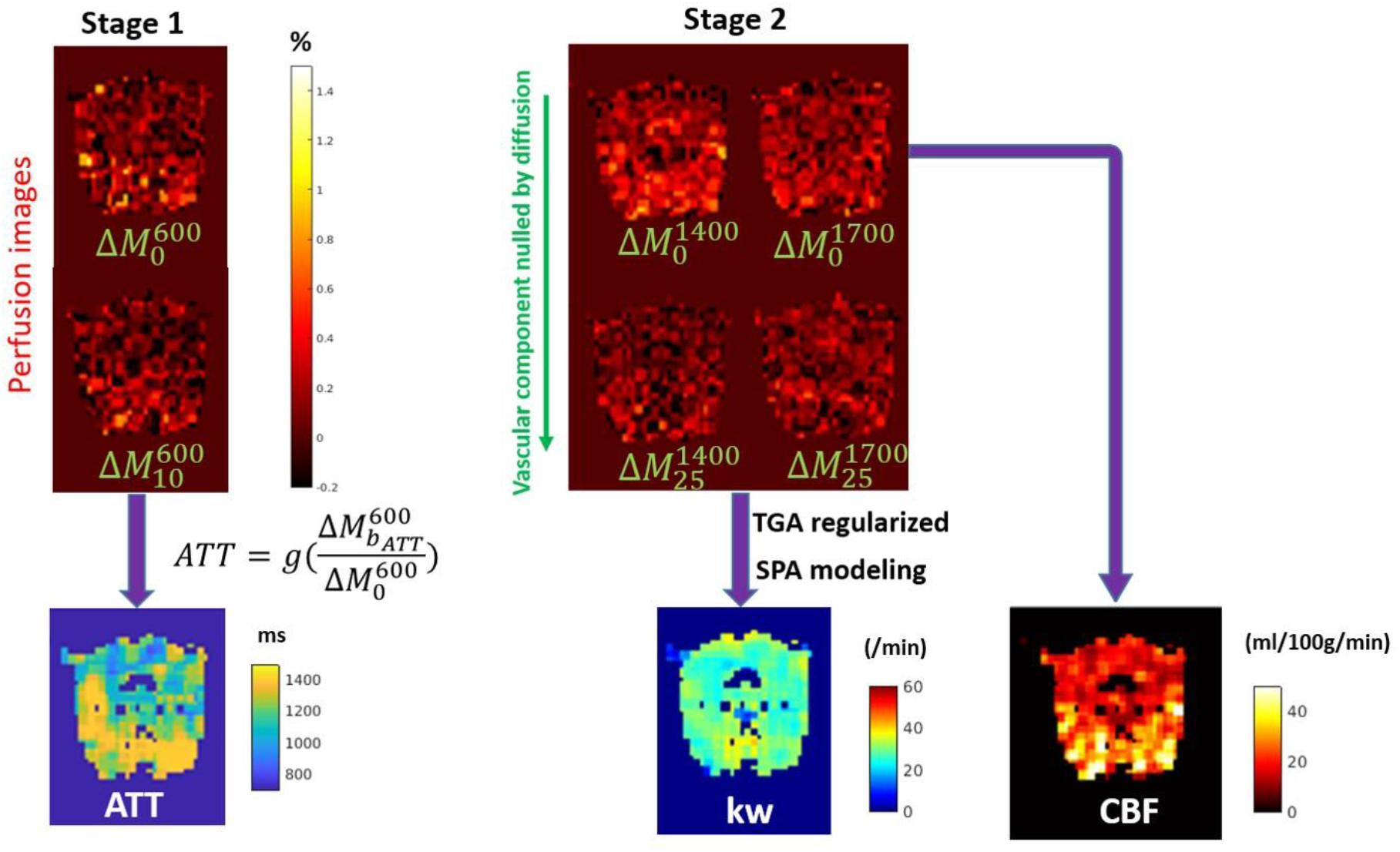
Two-stage estimation of arterial transit time (ATT), water‐exchange rate (*K*_*w*_), and cerebral blood flow (CBF). ATT is estimated via FEAST using the ratio of diffusion-weighted (*b*_*ATT*_ = 10 s/mm^2^) to non-weighted ASL signal at post-labeling delay (PLD) = 600 ms. *K*_*w*_ is derived from the two-compartment single-pass approximation model fitted to DP-pCASL data. CBF is calculated from 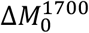 (b = 0 s/mm^2^, PLD = 1700 ms) without diffusion preparation. Here, 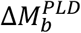 denotes the ASL difference signal with diffusion weighting b and delay PLD.

For localized quantitative analysis, specific regions of interest (ROIs) were defined on the single axial slice intersecting the dorsal striatum, selected from the limited field-of-view ASL acquisition. The planned target ROI, serving as the primary anatomical reference, was manually delineated directly on the parametric maps. Each ROI covered a 2 × 2 × 1 voxel volume (4.4 × 4.4 × 5 mm^3^), centered on the dorsal caudate nucleus as identified on the corresponding T2-weighted structural scan acquired at the same slice plane. The control ROI was defined by mirroring this planned target ROI to the contralateral hemisphere. To independently validate the targeting accuracy and characterize the physiological extent of the functional opening, a complementary Gd-defined target ROI was generated through a multi-step image processing pipeline. Difference maps were generated by subtracting pre-contrast T1-weighted images from post-contrast images. A signal intensity threshold was then applied (defined as the mean intensity of the difference map plus 2 × standard deviation) to identify enhancement. We manually removed artifacts arising from hyper-intense vessels and ventricular spaces to isolate tissue-specific extravasation, and the resulting mask was smoothed with a Gaussian kernel (σ = 1) to ensure spatial continuity. To account for inter-scan physiological variations, voxel-wise *K*_*w*_ values were normalized to the global 98th percentile value.

To verify the spatial consistency between the planned and Gd-defined target definitions, we assessed the distribution of water exchange rates across the different region types. The normalized voxel-wise values were compared across the control, planned target, and Gd-defined target groups using a one-way analysis of variance (ANOVA) followed by Welch’s t-tests for pairwise comparisons.

Comprehensive assessment of the FUS-BBBO effect was performed at the group level using permutation testing. For global physiological changes, whole-brain average values of CBF, ATT, and *K*_*w*_ were compared between non-BBBO (n = 6) and BBBO (n = 7) sessions using two-tailed independent-samples permutation tests in MATLAB R2023a (The MathWorks, Natick, MA, USA). Concurrently, the specific local treatment effect was evaluated by comparing the mean parameter values within the planned target ROI (BBBO) against the contralateral ROI (control) using paired-sample permutation testing. All permutation analyses utilized 10,000 iterations to determine statistical significance, with mean difference (Δ), p-value, and Hedges’ g reported. All co-registration and ROI operations were performed in FSL v6.0 (FMRIB Software Library, University of Oxford, Oxford, UK) (Jenkinson et al., 2012), with statistical significance defined at p < 0.05.

## Results

### Mapping of CBF, ATT, and K_w_ and its relation to contrast-enhanced T1

We first tested the effect of FUS-BBBO on hemodynamics and BBB water exchange. **Figure 4A** shows voxel‐wise maps of CBF, ATT, and BBB *K*_*w*_ in the axial plane through the dorsal striatum for two representative BBBO sessions in NHPs T and R. Both sessions targeted the right caudate and were acquired immediately after sonication with microbubbles. The CBF maps illustrate the baseline perfusion pattern throughout the striatum, with no systematic ipsilateral-to-contralateral differences attributable to BBBO. ATT maps display considerable variability across individual sessions, with average ATT values ranged from 918 to 1288 ms; however, there is no clear spatial correspondence between ATT heterogeneity and the BBBO locus. By contrast, *K*_*w*_ maps reveal pronounced, spatially restricted reductions in water-exchange rate at or immediately adjacent to the sonication site (white arrows), highlighting the unique sensitivity of *K*_*w*_ to focal BBB permeability changes. Furthermore, visual inspection of the raw EPI and structural T1-weighted images revealed no signs of macroscopic hemorrhage or edema in the sonicated regions, suggesting that the observed changes in water exchange rates were not confounded by tissue lesions.

**Figure 4.**
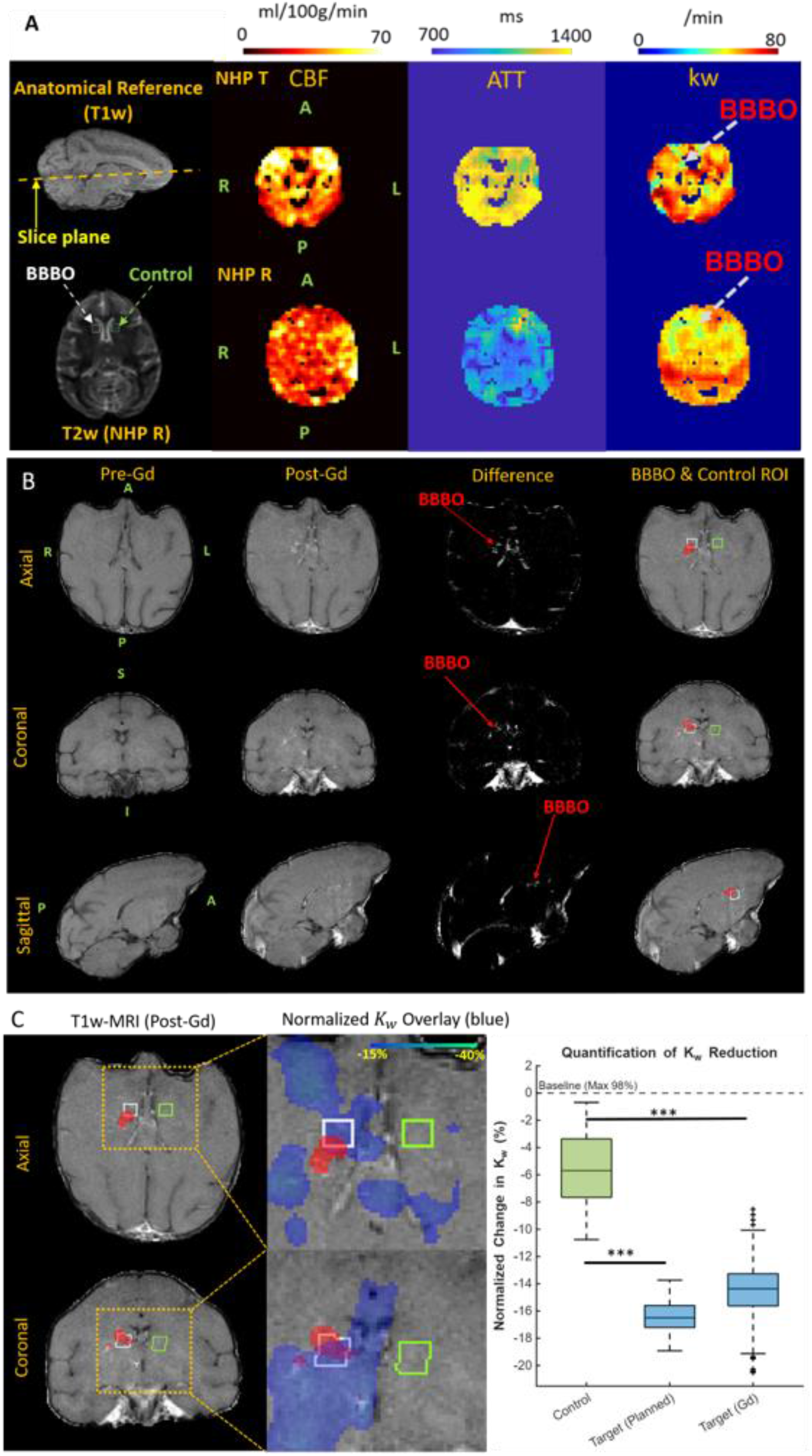
Spatial specificity of FUS-induced BBBO and cross-validation of *K*_*w*_ changes. (A) Representative parametric maps from NHPs R and T following FUS targeting the right caudate. Left column (Anatomical Reference) shows sagittal scouts indicating the ASL slice (yellow dashed line) and axial T2 images with planned target and contralateral control ROIs (square outlines). Subsequent columns show CBF, ATT, and *K*_*w*_ maps. Arrows and “BBBO” labels indicate focal *K*_*w*_ reduction within the sonicated region. (B) Anatomical verification. The Gd-defined target ROI (red mask) was generated from the difference between post- and pre-contrast T1 images, followed by thresholding, artifact removal, and smoothing. This overlay illustrates the spatial relationship between Planned Target (white box), Gd-defined Target (red mask), and Control (green box) ROIs. (C) Functional-anatomical concordance. Left: Zoomed overlay of *K*_*w*_ on T1 demonstrates specific *K*_*w*_ reduction (blue) within the Gd-enhanced BBBO region. Right: Quantitative comparison of normalized voxel-wise *K*_*w*_ across Control, Planned, and Gd-defined targets. ANOVA followed by Welch’s t-tests, reveals significant reductions in both targets vs. control, with highly consistent distributions between planned and Gd-defined targets (p < 0.001).

The spatial specificity of the observed *K*_*w*_ changes was verified by cross-referencing the functional maps with anatomical BBB opening. Figure 4B presents the multi-planar anatomical verification for a representative session, where comparison between the pre- and post-contrast T1-weighted images revealed a distinct, focal hyperintense region in the dorsal striatum. This confirmed successful contrast agent extravasation. The segmentation analysis demonstrated that the contrast-enhanced region (red mask) aligned precisely with the manually defined planned target ROI (white box), while the contralateral control ROI (green box) remained devoid of enhancement. To further validate *K*_*w*_ as a potential biomarker, we superimposed the *K*_*w*_ maps onto the high-resolution T1-weighted images. As illustrated in the zoomed-in panels of Figure 4C, the region of reduced *K*_*w*_ (blue overlay) exhibited excellent spatial concordance with the Gd-enhancement pattern. The reduction in water exchange rate was strictly confined to the sonicated area, establishing a direct link between BBB disruption and the local decrease in *K*_*w*_.

Quantitative analysis further confirmed a robust reduction in water exchange rates within the targeted regions. The control ROI exhibited a minor baseline deviation of -5.52% ± 2.62% relative to the global maximum, reflecting normal tissue heterogeneity. In contrast, the primary planned target ROI showed a substantial reduction of -16.39% ± 1.09%. Importantly, the complementary analysis using the Gd-defined ROI yielded a highly consistent reduction of -14.47% ± 1.68%. Statistical analysis revealed a significant difference between groups (one-way ANOVA, p < 0.001). Post-hoc tests confirmed that both the planned target and Gd-defined target groups differed significantly from the control group (p < 0.001), with large effect sizes (Cohen’s d values are 5.42 and 4.09 respectively). The statistical consistency between the geometrically planned target and the functionally defined Gd-mask confirms that the observed *K*_*w*_ reduction is a robust physiological effect, independent of the specific ROI definition method employed.

### Group-level comparison of whole-brain averages

To assess the reproducibility of BBBO effects, whole-brain CBF, ATT, and *K*_*w*_ were compared across four NHPs (non-BBBO: 6 sessions; BBBO: 7 sessions) via non-parametric permutation tests (**Figure 5A**). Whole-brain CBF showed no significant change following BBBO, with median (IQR) values of 31.3 (22.5–36.5) ml · 100 g^−1^ · min^−1^ in non-BBBO versus 29.3 (22.2–33.0) ml · 100 g^−1^ · min^−1^ in BBBO; permutation testing (10 000 iterations) yielded p = 0.64, a mean Δ = -3.0 ml · 100 g^−1^ · min^−1^, and Hedges’ g = -0.25. Whole-brain ATT shifted from a median (IQR) of 1204 (931–1264) ms in non-BBBO to 989 (930–1042) ms in BBBO; despite this apparent decrease, permutation testing (10 000 iterations) indicated no significant effect (p = 0.22; mean Δ = -106.9 ms; Hedges’ g = -0.65). In contrast, whole-brain *K*_*w*_ increased significantly after BBBO, with median (IQR) values of 49.8 (46.3 - 55.9) min^−1^ in non-BBBO versus 59.4 (56.6 - 66.3) min^−1^ in BBBO; permutation testing (10, 000 iterations) yielded p < 0.01, a mean Δ = 13.6 min^−1^ and Hedges’ g = 1.46, indicating a diffuse enhancement of water-exchange rate across the brain.

**Figure 5.**
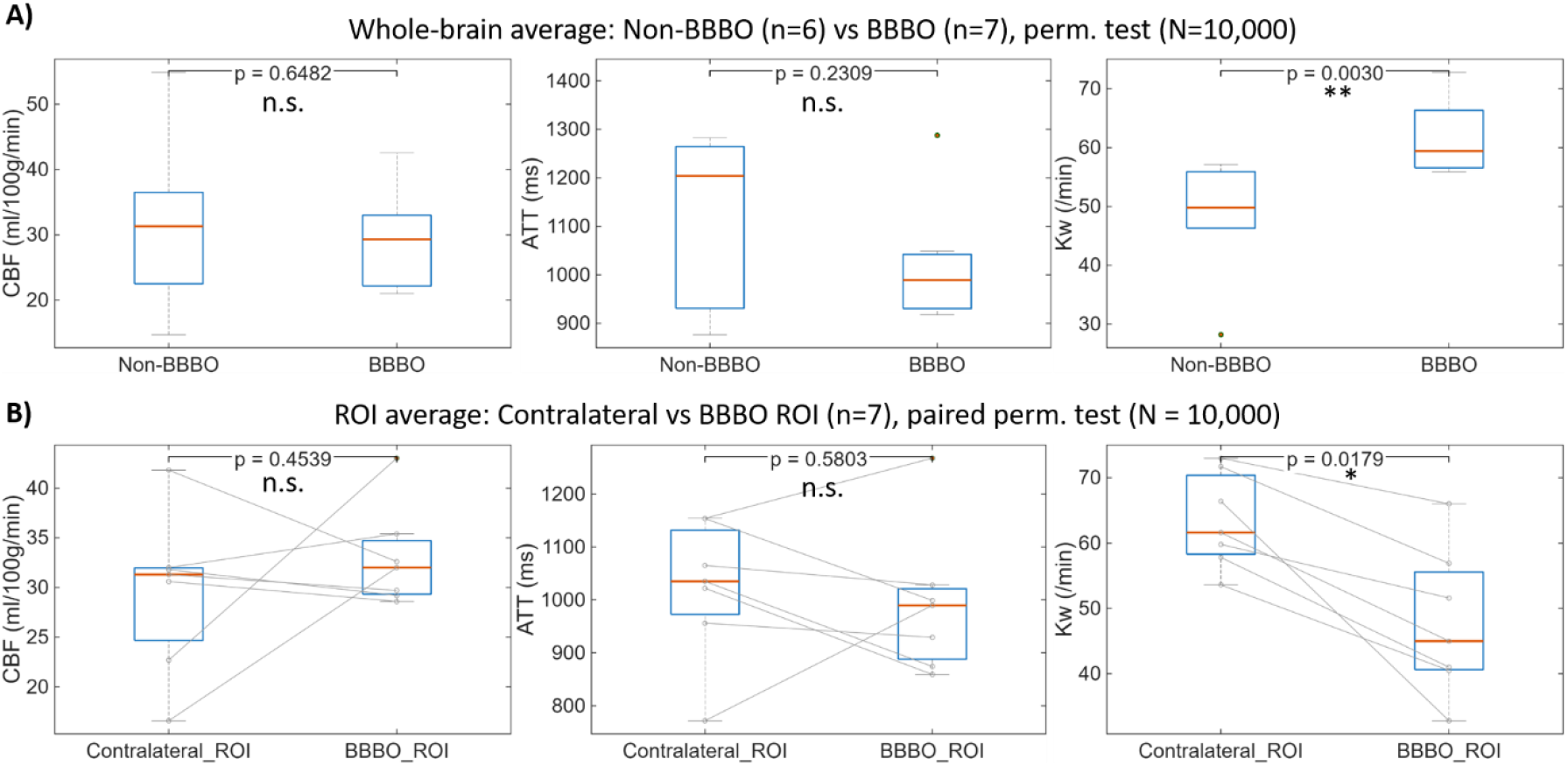
A) Whole-brain averages of cerebral blood flow (CBF), arterial transit time (ATT), and BBB water exchange rate (*K*_*w*_) in non-BBBO (n = 6 sessions, 4 NHPs) and FUS-BBBO conditions (n = 7 sessions, 4 NHPs). Permutation tests (N = 10,000) showed no significant differences in CBF and ATT (p > 0.05), but a significant increase in whole-brain *K*_*w*_ following BBBO (p < 0.01). B) ROI analysis comparing the BBBO site (caudate) and the contralateral control region within the same sessions revealed a significant reduction in *K*_*w*_ at the BBBO site (p < 0.05, paired permutation test), while CBF and ATT showed no significant differences (p>0.05). All box plots display the median, 25th and 75th percentiles (box edges), and individual outliers.

### ROI analysis between BBBO site and contralateral control

To assess focal effects of BBBO, CBF, ATT, and *K*_*w*_ were compared within the caudate ROI (planned target) at the BBBO site versus the mirrored contralateral control (n = 7 sessions) using two-tailed paired permutation tests (10,000 iterations; **Figure 5B**). CBF showed no significant change following BBBO, with median values of 31.3 (24.7-32.0) ml · 100 g^−1^ · min^−1^ in the contralateral control versus 32.0 (29.3 – 34.7) ml · 100 g^−1^ · min^−1^ in the BBBO ROI; permutation testing yielded p = 0.46, a mean Δ = 3.4 ml · 100 g^−1^ · min^−1^, and Hedges’ g = 0.27. ATT likewise showed no significant change, with median values of 1034.9 (972.5 – 1131.5) ms in control versus 989.6 (887.9 - 1120.9) ms in BBBO (p = 0.58; mean Δ = -30 ms; Hedges’ g = -0.17). *K*_*w*_ decreased significantly at the BBBO site, with median values of 61.6 (58.3 – 70.4) min^−1^ in control versus 45.0 (40.6 – 55.6) min^−1^ in BBBO; permutation testing yielded p < 0.05, a mean Δ = -15.7 min^−1^ and Hedges’ g = -1.51, indicating a localized suppression of water-exchange rate at the sonication locus.

## Discussion

This study demonstrates that FUS combined with microbubbles significantly reduces BBB *K*_*w*_ in the dorsal striatum of non-human primates. By contrast, our between-group analysis showed that whole-brain *K*_*w*_ was significantly higher in BBBO cases than in non-BBBO controls. The local reduction in *K*_*w*_ is consistent with a recent rodent study by Tiwari et al. (Tiwari et al., 2025), which used diffusion-weighted ASL MRI to assess BBB water exchange following chemically induced disruption and reported significantly reduced *K*_*w*_ in the affected hemisphere. Similar reductions in *K*_*w*_ have also been reported in individuals with neurological conditions such as aging, Alzheimer’s disease, and cerebrovascular disorders, where impaired BBB function has been frequently reported. (Chen et al., 2025; Palomares et al., 2015; Uchida et al., 2023; Wengler et al., 2020). Together, these converging results suggest that reduced *K*_*w*_ may serve as a common physiological signature of compromised BBB function, whether induced by external interventions or underlying disease, and support its application in evaluating the impact of FUS-BBBO.

The focal reduction in BBB *K*_*w*_ observed after FUS-BBBO suggests underlying cellular and molecular adaptations in response to barrier disruption. Aquaporin-4 (AQP4) water channels, primarily located on astrocytic endfeet, are key regulators of cerebral water homeostasis and glymphatic clearance (Mestre et al., 2018; Papadopoulos and Verkman, 2013). FUS-induced loosening or disruption of tight junctions may activate signaling pathways that alter AQP4 expression or polarization—potentially as a protective mechanism to regulate transvascular water flux and limit vasogenic edema (Cong and Kong, 2020). Similar regulatory responses have been reported in neuropathological conditions associated with BBB dysfunction, such as Alzheimer’s disease, stroke, and traumatic brain injury (Hirt et al., 2017; Tang et al., 2014; Zeppenfeld et al., 2017). Therefore, a reduction in *K*_*w*_ at the FUS-sonicated area may reflect an active homeostatic adjustment rather than a passive increase in permeability or leakage.

Moreover, our studies align with a growing consensus that BBB permeability is a dynamic, multifactorial phenomenon rather than a mere consequence of tight-junction disruption (Daneman and Prat, 2015; Sweeney et al., 2018). Endothelial remodeling, including changes in cell shape, junctional protein expression and transporter activity, pericyte detachment or loss, and astrocyte endfoot reactivity all modulate paracellular and transcellular flux (Abbott et al., 2006; Armulik et al., 2011). In parallel, neuroinflammatory mediators (e.g. IL-1β, TNF-α, matrix metalloproteinases, reactive oxygen species) and glymphatic flow disturbances each contribute to net water and solute exchange across the barrier (Kovacs et al., 2017; Mestre et al., 2018). By capturing the rapid FUS-induced changes in BBB *K*_*w*_ of non-human primates, DP-pCASL provides a single quantitative measure that integrates the net effects of endothelial remodeling, astrocyte reactivity, and inflammatory signaling on BBB water exchange (Kovacs et al., 2017; Shao et al., 2019).

In this study, we also observed a significant increase in the whole-brain *K*_*w*_ following FUS-BBBO targeted to the caudate nucleus. While the underlying mechanism remains unclear, a recent study has linked elevated whole brain *K*_*w*_ to improved executive function in older adults (Pappas et al., 2024). Although our study did not directly assess cognitive function after BBBO, earlier work has shown that FUS-BBBO can enhance visual behavior task performance in non-human primates by improving both accuracy and response speed, suggesting a potential neuromodulatory effect accompanying BBBO (Chu et al., 2015; Downs et al., 2017). Furthermore, recent work has demonstrated that FUS-BBBO in the dorsal striatum can activate broader brain networks including the default mode network and frontotemporal network (Liu et al., 2023). These findings raise the possibility that global *K*_*w*_ increases may reflect broader functional shifts in neurovascular regulation or fluid transport pathways beyond the sonicated target.

The observed *K*_*w*_ values in NHPs (median whole-brain average are 49.8 min^−1^ and 59.4 min^−1^ for non-BBBO and BBBO conditions respectively) are lower than those typically reported in awake humans (∼100-120 min^−1^). This discrepancy likely arises from the systemic effects of isoflurane anesthesia, which is known to alter cerebral blood flow, metabolism, and neurovascular coupling, as well as the use of different anesthetic protocols in NHP versus awake human studies (Li et al., 2014; Slupe and Kirsch, 2018; Zhang, 2022). In addition, ASL-based estimates of the BBB water exchange rate *K*_*w*_ have previously been shown to vary across species and experimental conditions, with animal studies often reporting different values with those in humans and attributing these differences to species-dependent capillary surface area, anesthesia, and modeling choices (Shao et al., 2023; Tiwari et al., 2017). Finally, imaging of the smaller NHP brain is more susceptible to partial volume effects compared to human studies, presenting a challenge for the precise quantification of ASL-derived kinetic parameters like *K*_*w*_ (Asllani et al., 2008; Chappell et al., 2021). However, to the best of our knowledge, this is the first study to quantify BBB water exchange rates in non-human primates using DP-pCASL. Therefore, these findings provide a critical baseline for future preclinical investigations into BBB permeability dynamics.

This study also investigates the potential of DP-pCASL as a non-invasive, contrast-free imaging biomarker for monitoring FUS-induced BBB permeability. Currently, dynamic contrast enhanced MRI (DCE-MRI) is widely used to evaluate BBB integrity by measuring the leakage of contrast agents (CA) into the tissue’s interstitial space, indicated by increased T1-weighted signal intensity. Using pharmacokinetic model computations, the volume transfers constant *K*^*trans*^ serves as a surrogate measure of BBB permeability (Sourbron and Buckley, 2013). However, the safety of exogenous parametric CA (i.e. Gadolinium) has become a concern for human health (Gulani et al., 2017; Harvey et al., 2020). Recent research has focused on non-Gd-based methods for imaging the BBB integrity, including the use of D-glucose as a CA for chemical exchange saturation transfer (CEST) MRI (Xu et al., 2015), and arterial spin labeling (ASL) for measuring water exchange as a marker for non-contrast detection of BBB impairment (Li et al., 2022). Diffusion tensor imaging (DTI) has been investigated as a non-contrast method for BBB opening detection, linking changes in fractional anisotropy to BBB permeability (Karakatsani et al., 2020). However, DTI provides no direct hemodynamic or perfusion metrics, such as cerebral blood flow or volume, that are more physiologically tied to barrier function (Le Bihan and Iima, 2015). In contrast, DP-pCASL specifically quantifies BBB water exchange rates by differentiating labeled water in vascular and extravascular compartments using diffusion weighting and kinetic modeling (Shao et al., 2019; St. Lawrence et al., 2012). This provides a direct and physiologically meaningful measurement of BBB permeability to water, an endogenous tracer with small molecular size and ubiquitous presence.

This study did not reveal sustained or statistically significant changes in CBF or ATT at either the whole-brain level or within the BBBO ROI. Although Labriji et al. (2023) reported a 29.6 ± 15.1 % decrease in CBF and prolonged ATT following BBBO in rats (Labriji et al., 2023), our NHP data showed only modest, non-significant fluctuations (whole-brain CBF Δ = –3.0 mL·100 g^−1^·min^−1^; p = 0.64; whole-brain ATT Δ = –106.9 ms; p = 0.22). These discrepancies may reflect differences in BBBO targeting and volume, species differences (earlier study in rodents versus our work in non-human primates), and the limited longitudinal sampling in our current protocol, which may have missed the critical timeline of CBF drops (Rigollet et al., 2024). Additionally, isoflurane anesthesia, a potent cerebral vasodilator, likely alters vascular responsiveness during these experiments, thereby masking subtle changes induced by BBBO and further limiting CBF and ATT as reliable, noninvasive biomarkers of BBBO in NHPs (Slupe and Kirsch, 2018).

It is worth noting that the observed whole-brain CBF range of 14.7 - 54.9 ml/100g/min is broadly consistent with the physiological range reported in a recent NHP pCASL study by Johnson et al., which documented a baseline whole-brain CBF range of 21.47 - 77.23 ml/100g/min (Johnson et al., 2024). While the median values in present study (31.3 ml/100g/min and 29.3 ml/100g/min for non-BBBO and BBBO conditions respectively) are lower than their reported mean (∼51.5 ml/100g/min), this difference can be contributed to the dose-dependent vasodilator effect of isoflurane (Li et al., 2014). Specifically, a lower maintenance concentration was used in the current protocol (0.8–1.1%) compared to the 1.5% employed by Johnson et al. Notably, the measurements align closely with gold-standard PET values obtained under propofol anesthesia which reported a whole brain CBF range of 23 - 42 ml/100g/min (Kudomi et al., 2005). Furthermore, while earlier studies using continuous ASL reported much higher CBF (∼104 ml/100g/min in grey matter)(Zhang et al., 2007), those elevations may be attributable to methodological differences inherent to continuous labeling techniques coupled with deeper anesthesia.

The study has several limitations. First, the spatial resolution is primarily constrained by the signal-to-noise ratio (SNR), which is influenced by acquisition time and coil sensitivity. Consequently, scattered low *K*_*w*_ regions located near the brain edges, ventricles, and sulci likely reflect inherent signal fluctuations and partial volume effects. However, within the brain parenchyma, the region of reduced *K*_*w*_ extended beyond the boundaries of Gd-enhanced region, Given the differential sensitivity between water (∼18 Da) and gadolinium-based contrast agents (∼550 Da), it is biologically plausible that FUS induces functional alterations in water dynamics in peripheral areas where the barrier is modulated but not physically disrupted enough to leak contrast agent. Additionally, we cannot rule out that some remote changes represent genuine physiological modulation coupled with functional network shifts (Liu et al., 2023), although characterizing these distinct physiological fluctuations remains a subject for future investigation. To overcome these current resolution limits, recent motion-compensated, multi-delay diffusion-weighted ASL with isotropic 3.5 mm^3^ resolution in humans (Shao et al., 2023) may substantially improve both sensitivity and resolution.

Second, DP-pCASL was acquired at approximately 45-60 min post-sonication, capturing only offline effects rather than the full opening–recovery time course. Longitudinal, multi-time point imaging will be essential to elucidate how FUS-BBBO dynamically alters cerebral blood flow and water-exchange metrics over time (Labriji et al., 2023; Rigollet et al., 2024; Sun et al., 2015). Third, potential limitations related to repeated isoflurane anesthesia and the absolute Kw value must be carefully considered. While *K*_*w*_ is highly sensitive to changes in BBB permeability, the absolute *K*_*w*_ values reported in this study, which were obtained under isoflurane anesthesia, may not perfectly reflect the values of an awake NHP due to the systemic effects of volatile agents (Slupe and Kirsch, 2018). However, in the context of this within-subject design focusing on changes, the stability of CBF and ATT (systemic and anesthesia-sensitive metrics) across sessions strongly suggests that the observed increase in *K*_*w*_ is not caused by a general cumulative systemic effect (e.g., gradual shift in vascular tone or ETCO2 drift). Instead, the change in *K*_*w*_ is more likely attributed to the targeted BBBO procedure itself and its interaction with global brain networks.

Fourth, our single-pass approximation kinetic model simplifies the microvascular environment and does not fully capture transcellular water movement through AQP4 channels or intra-voxel transit-time heterogeneity. While we accounted for voxel-wise arterial transit time (ATT) in this study, further refinements could incorporate mechanisms suggested by recent multi-echo time ASL studies, such as using varying echo times to probe AQP4-associated transcellular flux (Ohene et al., 2019), or modeling intra-voxel transit-time dispersion to improve estimation accuracy (Mahroo et al., 2021). Incorporating these advanced models and validating the resulting *K*_*w*_ values against dynamic contrast-enhanced MRI–derived *K*^*trans*^ measurements (Hývlová et al., 2025; Tiwari et al., 2025; Vlachos et al., 2010) could enable precise mapping of how FUS affects both the timing and spatial distribution of transcellular and paracellular water exchange.

In conclusion, our findings demonstrate that FUS-mediated BBB opening results in decreased *K*_*w*_ at the focal site alongside a global increase in whole-brain *K*_*w*_, revealing complex physiological adaptations that can be noninvasively monitored using DP-pCASL. This technique offers a safe, repeatable, and sensitive tool for detecting and tracking BBB permeability changes, with strong potential for both preclinical research and clinical application in neurotherapeutics.

## Credit author statement

D.L. and V.P.F. designed the study, collective and analyzed the data and wrote the manuscript. X.S. and D.J.J.W. provided critical support for implementing the DP-pCASL sequences and processing the data. R.L. configured the MRI sequence and assisted with data collection. F.M. and S.S. helped with study design, device calibration, and data collection. E.E.K. provided focused ultrasound expertise. V.P.F. and E.E.K. supervised the study.

## Acknowledgements

The authors wish to acknowledge the ZI-ICM team for the setup and anesthesia procedures during MRI imaging, and ZI-MRI team for the MRI sequence setup and data collection. The work was supported by NIH R01 MH133020 (V.P. Ferrera and E.E. Konofagou), R01 NS134712 (D.J.J. Wang) and BBRF Young Investigator grant 31298 (D. Liu).

